# Human telomere length is chromosome specific and conserved across individuals

**DOI:** 10.1101/2023.12.21.572870

**Authors:** Kayarash Karimian, Aljona Groot, Vienna Huso, Ramin Kahidi, Kar-Tong Tan, Samantha Sholes, Rebecca Keener, John F. McDyer, Jonathan K. Alder, Heng Li, Andreas Rechtsteiner, Carol W. Greider

## Abstract

Short telomeres cause age-related disease and long telomeres predispose to cancer; however, the mechanisms regulating telomere length are unclear. To probe these mechanisms, we developed a nanopore sequencing method, Telomere Profiling, that is easy to implement, precise, and cost effective with broad applications in research and the clinic. We sequenced telomeres from individuals with short telomere syndromes and found similar telomere lengths to the clinical FlowFISH assay. We mapped telomere reads to specific chromosome end and identified both chromosome end-specific and haplotype-specific telomere length distributions. In the T2T HG002 genome, where the average telomere length is 5kb, we found a remarkable 6kb difference in lengths between some telomeres. Further, we found that specific chromosome ends were consistently shorter or longer than the average length across 147 individuals. The presence of conserved chromosome end-specific telomere lengths suggests there are new paradigms in telomere biology that are yet to be explored. Understanding the mechanisms regulating length will allow deeper insights into telomere biology that can lead to new approaches to disease.

## Introduction

Human health is profoundly affected by telomere length, yet the detailed mechanism of length regulation is poorly understood. Telomere length is maintained as an equilibrium distribution with constant shortening at each round of DNA replication, which is counterbalanced by *de novo* addition of new telomere repeats by the enzyme telomerase ^1^. Failure to maintain the length distribution leads to inherited Short Telomere Syndromes and age-related degenerative disease such pulmonary fibrosis, immunodeficiency, and bone marrow failure ^2^. Conversely long telomeres predispose people to cancer ^3^, and a cluster of mutations that increases telomerase activity is one of the most common mutational signatures in cancer ^4,5^.

Precisely how telomerase action maintains a length equilibrium is of great interest. The prevailing ‘protein counting’ model for length maintenance ^6,7^ proposes that proteins that bind telomeric TTAGGG repeats negatively regulate telomere elongation in *cis.* This is supported by evidence that telomerase stochastically elongates short telomeres more frequently than long telomeres ^8^. Together these studies propose that the length equilibrium is maintained by telomerase lengthening short telomeres to precisely counterbalance shortening of all telomeres. An implication of this model is that all telomeres will be regulated around a similar mean length distribution.

Methods for measuring telomere length have had significant influence on length regulation models. Southern blotting with a telomere repeat probe was first established to measure the length of all telomeres in a population of cells and revealed a heterogeneous distribution of lengths ^9^ ^10^. The distribution is difficult to quantitate, and absolute lengths differ significantly between labs ^11^. The protein counting model was based on data from Southern blots which explains the focus on the regulation of the distribution of lengths across all telomeres ^12–14^. The clinical FlowFISH assay ^15^ used for diagnosis of telomere diseases ^16–18^ is normalized to the median length of telomeres on a Southern blot. The fact that this method is robust and accurately identifies telomere mediated disease may imply to some that the global telomere length average is the biologically relevant measurement, when in fact it might not be.

Other methods such as qFISH allows measurement of individual telomeres by *in situ* hybridization to a fluorescent probe in a metaphase spread ^19^. qFISH experiments have suggested that telomeres on all chromosome arms are not globally regulated around a common length distribution ^20–22^, however this data in not yet reconciled with the prevailing protein counting model of length regulation. FlowFISH and qFISH are not accessible to researchers outside specialized telomere biology labs, highlighting the need for a reproducible, accurate, and accessible tool for telomere length measurements. Here we describe nanopore Telomere Profiling, which measures the length of each individual telomere in the cell at nucleotide resolution. Using this technique, we establish that indeed individual telomeres on specific chromosome ends are maintained around their own unique length distributions, and the length distributions can differ by more than 6kb. Telomere profiling represents a paradigm shift in telomere analysis and will enable exploration of entirely new areas of telomere biology.

## Results

To determine whether human telomeres are maintained around a common length distribution across all chromosome ends or if specific chromosome ends maintain their own unique length distributions, we developed the Telomere Profiling method to physically enrich telomeres and sequence the using Oxford Nanopore Technology (ONT) MinION for long read sequencing. We ligated the telomeric ends with a biotinylated oligonucleotide (TeloTag) that contains a multiplexing barcode and restriction enzyme sites. Following ligation, we pulled down the tagged telomeres with streptavidin beads and released them by restriction enzyme digestion prior to sequencing (Fig. 1a). To assess the enrichment efficiency, we prepared libraries from both telomere enriched and non-enriched samples and sequenced them on an ONT MinION. Enrichment recovered ∼17% of total telomere input (Extended Data Fig. 1. b, c), and resulted in a ∼3400-fold increase in sequenced telomeres (Extended Data Fig.1d and methods). We routinely multiplexed samples and generated ∼50,000 telomere reads per flow cell with an average fragment length (subtelomere + telomere) of ∼20kb. The cost per sample when multiplexing was approximately $80-140 (see methods).

**Fig.1.**
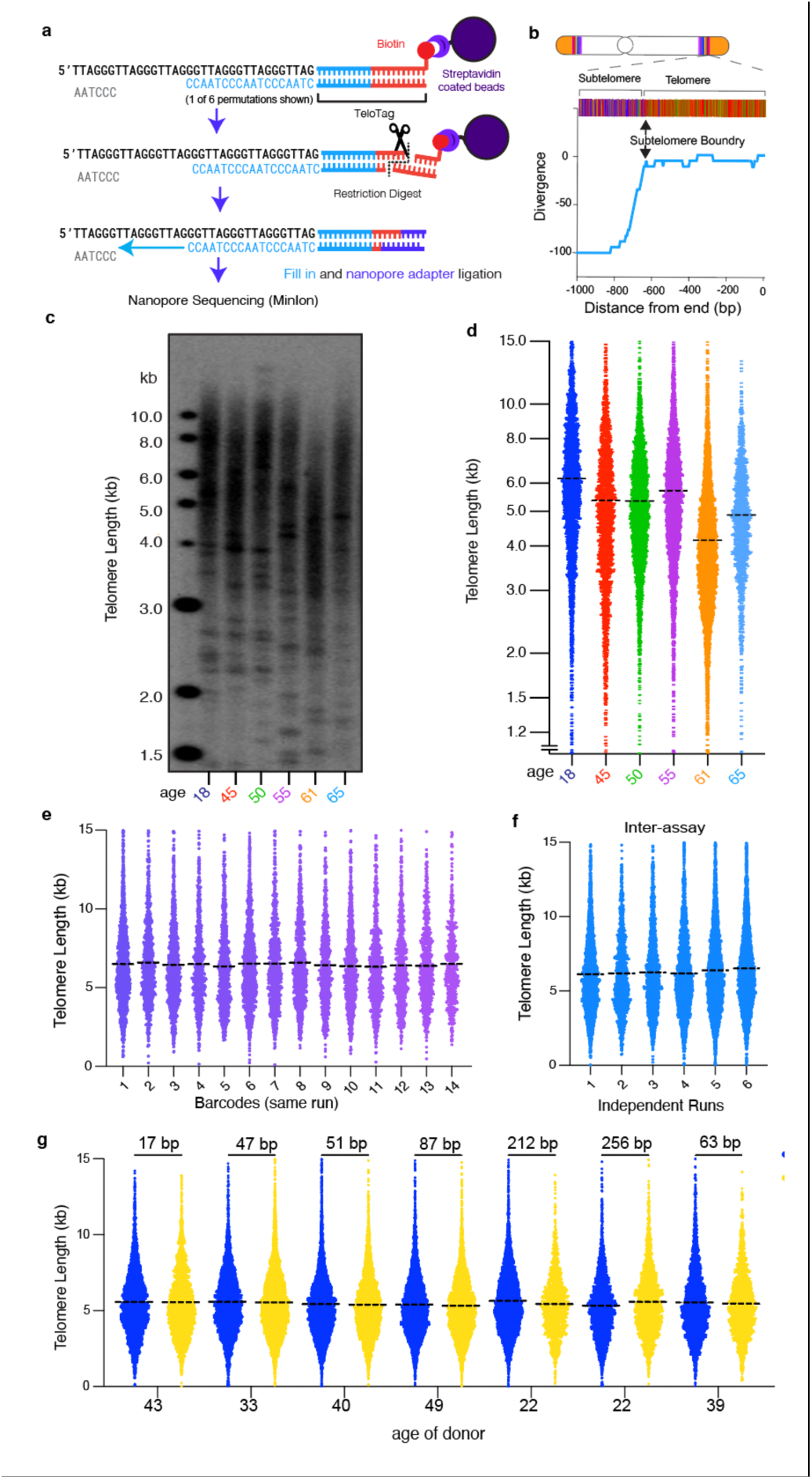
Nanopore telomere profiling is accurate and precise. **a** Schematic depicting nanopore telomere profiling enrichment strategy. Telomeres are tagged with a biotin adapter (TeloTag), enriched by streptavidin pull down, and sequenced. **b** Subtelomere boundary is identified using an algorithm that detects significant deviation from the telomere repeat pattern. **c.** Southern blot of telomere lengths from 6 individuals. **d**. Telomere length measured by Nanopore telomere profiling for the same individuals as in **c.** (total reads = 21,556). The dashed line represents the mean telomere length of each distribution. Each point represents a single telomere read. **e** Inter assay variability. Telomere length from a single donor was measured 14 times on a single flow cell (total reads = 13,256). **f.** Inter-assay variability: Telomere length measured from a single donor across 6 flow cells (total reads = 19,230) **g.** Telomere length profiles from the same samples generated in two different laboratories JH=Johns Hopkins (blue), SC=UC Santa Cruz (gold). The difference in telomere length in base pairs is shown at the top.

### Bioinformatic analysis of telomere length

We developed a bioinformatic pipeline to determine both “bulk length” (all telomeres) as well as chromosome end-specific telomere length. We used ONT Guppy for nucleotide base calling and filtered for reads containing telomere repeats (Methods). We initially determined telomere length using a method we used in yeast ^23^ that has been previously used in human cells ^24–26^. Reads are first mapped to a reference genome with a defined telomere/subtelomere boundary and telomere length is defined as the number of base pairs from the boundary to the end of the read (Methods). However, we found heterogeneity in subtelomere sequences among individuals, where sometimes the subtelomere was slightly longer or shorter than in the reference genome which caused overcalling or under calling of telomere lengths (Extended Data Fig. 2). To overcome this, we developed TeloNP, an algorithm to define the subtelomere boundary and measure telomere length directly from the nanopore sequencing reads while taking systematic nanopore basecalling errors ^27^ into account. TeloNP scans from the end of the read and defines the subtelomere boundary where it finds a sustained deviation from the expected telomeric sequence content (Fig. 1b and Extended Data Fig. 3) (see Methods). Telomere length was defined as the base pairs from the TeloTag to the subtelomere boundary identified by TeloNP in the telomere read.

To examine whether telomere length determined by TeloNP after Guppy base calling accurately represents the true length of the telomere repeats, we examined the electrical current signals from the flow cells. We developed TeloPeakCounter to count the repeated current peaks corresponding to (TTAGGG)n in telomere sequences and estimate telomere length (Extended Data Fig. 4a, b) (see methods). We found telomere length determined by TeloNP after Guppy base calling was in good agreement with length determined by TeloPeakCounter. We therefore adopted Guppy base calling for our analyses. We note the new ONT Dorado base caller (version 0.3.1) overestimated telomere length for G strand reads compared with both TeloPeakCounter and Guppy (Extended Data Fig. 4 c, d).

### Nanopore Telomere Profiling accurately and reproducibly reports telomere length

We performed Telomere Profiling by sequencing DNA from Blood or PBMCs (see methods) of 132 people ranging from 0 years to 91 years of age (Fig. 1d) and found agreement with telomere lengths on a Southern blot (Fig. 1c). To test reproducibility, we measured telomere length of DNA from one individual on the same flow cell (intra-assay) (Fig. 1e and Extended Data Fig. 1e) (coefficient of variation, CV 1.3%) or on different flow cells (inter-assay) Fig. 1f and Extended Data Fig. 1f (CV 2.4%). This low variability compares well with FlowFISH that has an inter-assay CV of 2.2%, and considerably outperformed the frequently used qPCR assay that has an inter-assay CV of 25.0% ^17^. In addition, we tested inter-lab variability by measuring telomere length of seven samples where the same DNA was enriched and sequenced by two different people in two different labs (Johns Hopkins and UCSC) and found highly reproducible results (mean difference of 104.7 bp with SEM of +/- 34 bp) (Fig. 1g). To determine whether any fragment length bias of nanopore sequencing could skew telomere length determination, we compared restriction enzyme cutting with a combination of *BamHI* and *EcoRI* which generates fragments ∼9 kb, or with *AsiSI* and *PvuI,* which generate fragments ∼25 kb. We found similar telomere lengths in these two samples (6,127 bp vs 6,188 bp) (Extended Data Fig. 5a, b) indicating fragment length in these size ranges did not have detectable bias on telomere length determination.

### Telomere profiling determines telomere shortening with age at nucleotide resolution

Telomere length is known to shorten with age ^17,28–31^, however previous methods could not measure telomere length at nucleotide resolution. To test the dynamic range of Telomere Profiling, we first applied nanopore Telomere Profiling to DNA samples of 11 individuals from 0-84 years of age and ordered the samples based on decreasing telomere length (Fig. 2a). We then did a Southern blot on the same DNA and found Telomere Profiling predicted the relative order of telomere lengths and captured the wide dynamic range of the Southern blot (Fig. 2a and b). Southern blotting does not measure the shortest telomeres because telomere repeats are required for probe hybridization on a Southern. We plotted the 1st, 10th and 50th percentile of telomere length as determined by Telomere Profiling and observed a decrease of the 50th and 10th percentile as the mean length shortened. However, the 1st percentile telomere length did not decrease suggesting there is a threshold length in PBMC’s of below which telomeres cannot be maintained (Fig. 2c).

**Fig. 2.**
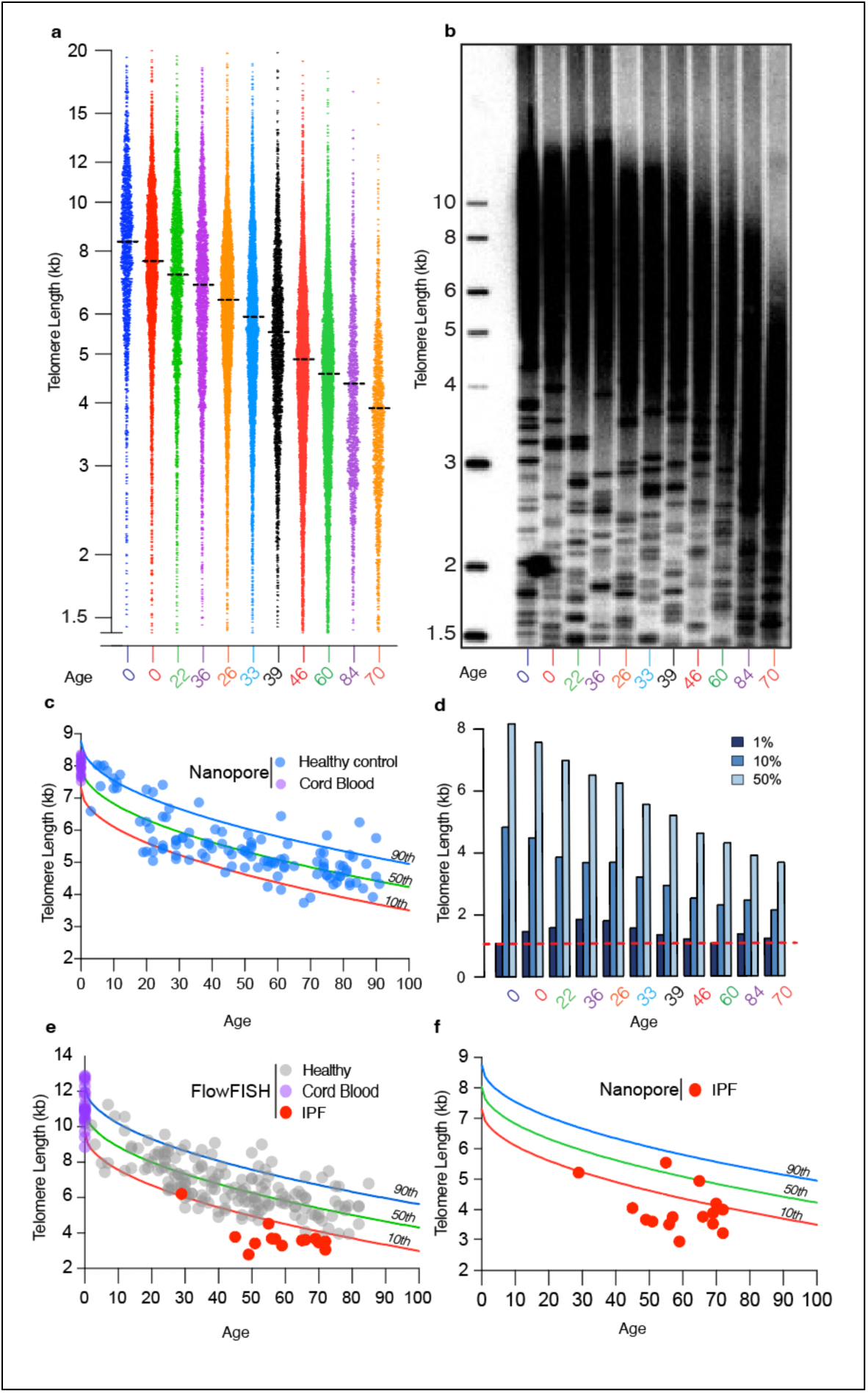
Nanopore telomere profiling identified population distribution of telomere shortening with age similar to FlowFISH. **a**. Nanopore telomere length profiles for 11 samples selected, age is noted at bottom. Each point is an individual read (total reads = 47624) **b.** Southern of same samples as **a.**, Age of individual noted at bottom **c**. The mean telomere length was determined for 132 individuals aged 0 to 91 (blue dots) (total reads = ∼920,000). Cord blood lengths are shown in purple. We calculated the population distribution and show confidence intervals for the 90th percentile (blue), the median (green line) and 10th percentiles (red) for telomere length in this population using parameters established previously for FlowFISH ^17^ **e**. Lymphocyte telomere length from FlowFISH data from Alder et al. (gray dots) and cord blood (purple dots). The lengths for 15 human subjects with IPF (red) were determined by FlowFISH (Methods) One point represents two individuals who have nearly identical length and are indistinguishable in the figure. **f**. Nanopore telomere length profiles from the same 15 subjects with short telomere syndrome shown in **e.** plotted against population distribution from **c** (total reads =32457).

To directly compare nanopore Telomere Profiling to FlowFISH, we conducted nanopore Telomere Profiling on whole blood and PBMCs (see methods) of 132 donors ranging from 0 to 90 years of age (Fig. 2d). Using the mean telomere length for each individual, we defined 90^th^, 50^th^ and 10^th^ intervals of telomere length at each age using the same statistical methods used for FlowFISH ^17^. While the shapes of the curves are very similar between Telomere Profiling and FlowFISH, the absolute lengths of the telomeres are longer for FlowFISH. Nanopore Telomere Profiling of cord blood showed a telomere length distribution with a mean of 7,986 +/- 245 bp (Fig. 2d) across 18 samples. This is shorter than the average cord blood telomere length for FlowFISH and with less variance (Fig. 2e) ^17^. Average cord blood telomere length estimates measured by FlowFISH vary from ∼18kb ^32^ to ∼9kb ^33^ to ∼11 kb ^17^. FlowFISH fluorescence signal is normalized to Southern blots, which includes some subtelomeric sequences and this may account for the longer telomere lengths of FlowFISH. Furthermore, Southern blot estimated telomere lengths are known to vary between laboratories ^11^. In contrast, nanopore Telomere Profiling offers a precise readout in base pairs that can be directly compared between laboratories.

To compare our method directly to FlowFISH, we sequenced 5μg of archived DNA from blinded samples of individuals previously diagnosed with Idiopathic pulmonary fibrosis (IPF) one of the Short Telomere Syndromes ^34^. Telomere profiling showed that bulk telomere length in most IPF samples were similar to the FlowFISH measurement (Fig. 2f). FlowFISH uses flow cytometry and can distinguish telomere lengths in specific cell types from whole blood samples ^35,36^ and some samples have discordant lymphocyte and granulocyte telomere lengths ^17,37,38^, Nanopore Telomere Profiling will report the average length from all cell types in the samples. Thus, while nanopore Telomere Profiling likely can be used for diagnosing Short Telomere Syndromes in the future, additional development such as isolation of specific cell types may help to capture heterogeneity of clinical samples.

### Human telomeres have chromosome end-specific length and haplotype-specific length differences

To determine whether humans have chromosome end-specific telomere length, we first examined telomeres from the diploid HG002 cell line for which a high-quality reference genome is available ^39^. Human subtelomeres contain many blocks of homology shared between different telomeres (paralogy blocks) ^24,40^. Simulation of long read data from CHM13 references genome showed that *minimap2* ^41,42^ can assign simulated reads to the correct telomere with high accuracy using 10kb of subtelomere sequence ^27^. We isolated DNA from HG002 cell line, sequenced the telomeres and mapped reads with an average total length of 16.4 kb (4.6 kb telomere repeats and 11.8 kb sub-telomeric sequence, on average) to the HG002 reference genome using *minimap2* using a customized filtering pipeline (methods). Seventy-seven chromosome ends passed our quality filters, and we found 66 ends had significant differences in length distribution from the grand mean (Fig. 3. a, b). In addition to chromosome end-specific lengths, we also found that some telomeres showed significant differences between the maternal and paternal haplotypes. In some cases, remarkably, there was more than 6kb difference in mean length, for example for chromosome 1p Maternal (1pM) and 1p Paternal (1pP). Thus, like in yeast ^23^, humans have chromosome end-specific telomere length distributions.

**Fig. 3.**
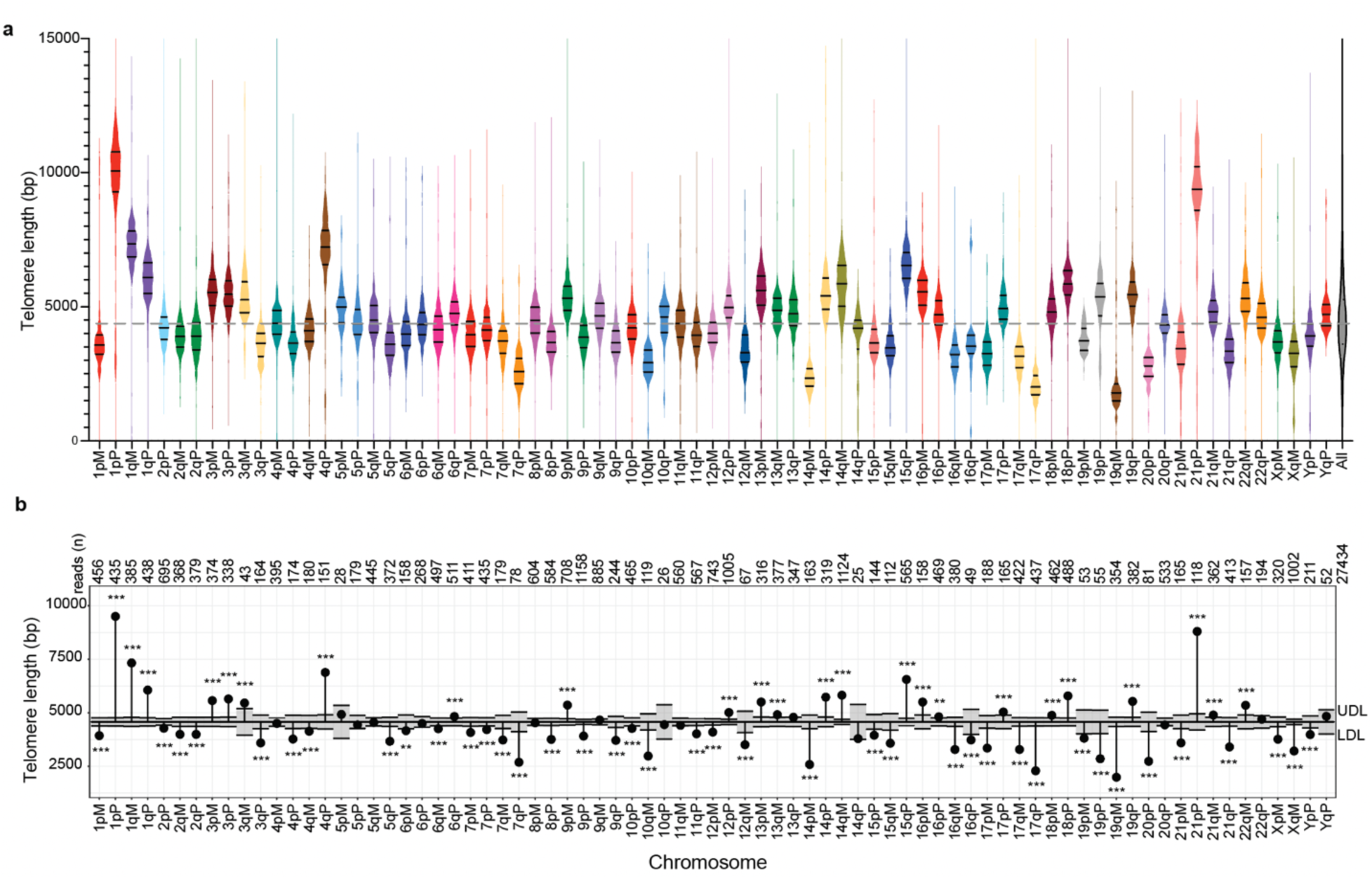
Chromosome end-specific telomere lengths. **a.** Violin plots of the distribution of telomere lengths for 77 telomeres from HG002 that mapped with confidence and passed our filters (see methods) Total reads n=27,433. Each end is labeled with the chromosome number and p for the short and q for long arms. The haplotypes for each chromosome end are labeled Maternal (M) and Paternal (P) and colored with the same colors for allelic pairs. The mean, 90th and 10th percentile for each distribution are shown with short horizontal black lines in each violin plot. The distribution of all telomeres lengths across all chromosomes ends is at the far right (All). The dashed line represents the grand mean of all telomeres **b**. Analysis of the means (ANOM) multiple contrast test of each telomere length distribution against the grand mean of all telomere lengths for data in **a.** The number of reads for each chromosome end is shown at the top. P-values were adjusted for multiple hypotheses testing using the Bonferroni method. Chromosome ends with length profiles reaching outside of the shaded gray region between the upper decision limit (UDL) and lower decision limit (LDL) are considered significantly different from the grand mean. (*) p ≤ 0.05, (**), p ≤ 0.01. (***), p ≤ 0.001; nonsignificant differences have no stars.

### Chromosome specific telomere lengths are conserved across individuals

To determine whether chromosome end-specific differences were conserved across a broad population, we used *minimap2* to map ∼920000 telomere reads from 147 individuals to the subtelomere sequences from the recently released pangenome containing 47 high quality T2T assemblies ^43^ and filtered for reads with >1kb of alignment, which resulted in ∼647,000 reads (see methods). We removed the acrocentric and X Y chromosome ends because the high rate of meiotic recombination between these ends across a population would not allow them to map uniquely ^44^.

*Minimap2* map quality score (mapq) is not optimized for mapping to the multiple genomes present in the pangenome as most reads have multiple near identical alignments and thus get low mapq. To establish if reads reproducibly mapped to the same subtelomere, we compared the pangenome alignment of reads to their alignment in three different high quality haploid reference T2T genomes CHM13, HG002 maternal and HG002 paternal. Of the ∼647,000 reads that aligned to the pangenome ∼350,000 mapped to the T2T references with a mapq of 60. We compared the fraction of reads that were mapped to a given chromosome end in the pangenome (column) to where they mapped in the respective T2T haploid genome (rows) in a matrix heatmap (Fig. 4 a, b, c). The diagonal indicates the fraction of reads mapping to a chromosome in the pangenome that map to the same chromosome end in the respective haploid genomes. 87% of the filtered reads mapped to the same chromosome end in the pangenome and CHM13, 90% in the pangenome and HG002 maternal and 88% in the pangenome and HG002 paternal. We also quantified the percent of reads that mapped to the same chromosome end in the pangenome and all three haploid reference genomes (Fig. 4d and Extended Data Fig. 5 a, b, c). For 33 of the 39 chromosomes ends, 100-60% of the reads mapping to a given chromosome end in the pangenome mapped to the same end and all three haploid genomes. Six chromosome ends had between 10-20% of reads map to the same chromosome end in the pangenome and all three haploid genomes. When we added back the acrocentric chromosomes, we found 0 reads mapped back to the same chromosome end in all three references (Extended Data Fig 5d), as expected for reads that map to several different chromosome ends across a population. Together this data suggests that the reads we found mapping to a certain pangenome chromosome map with high confidence.

**Fig. 4.**
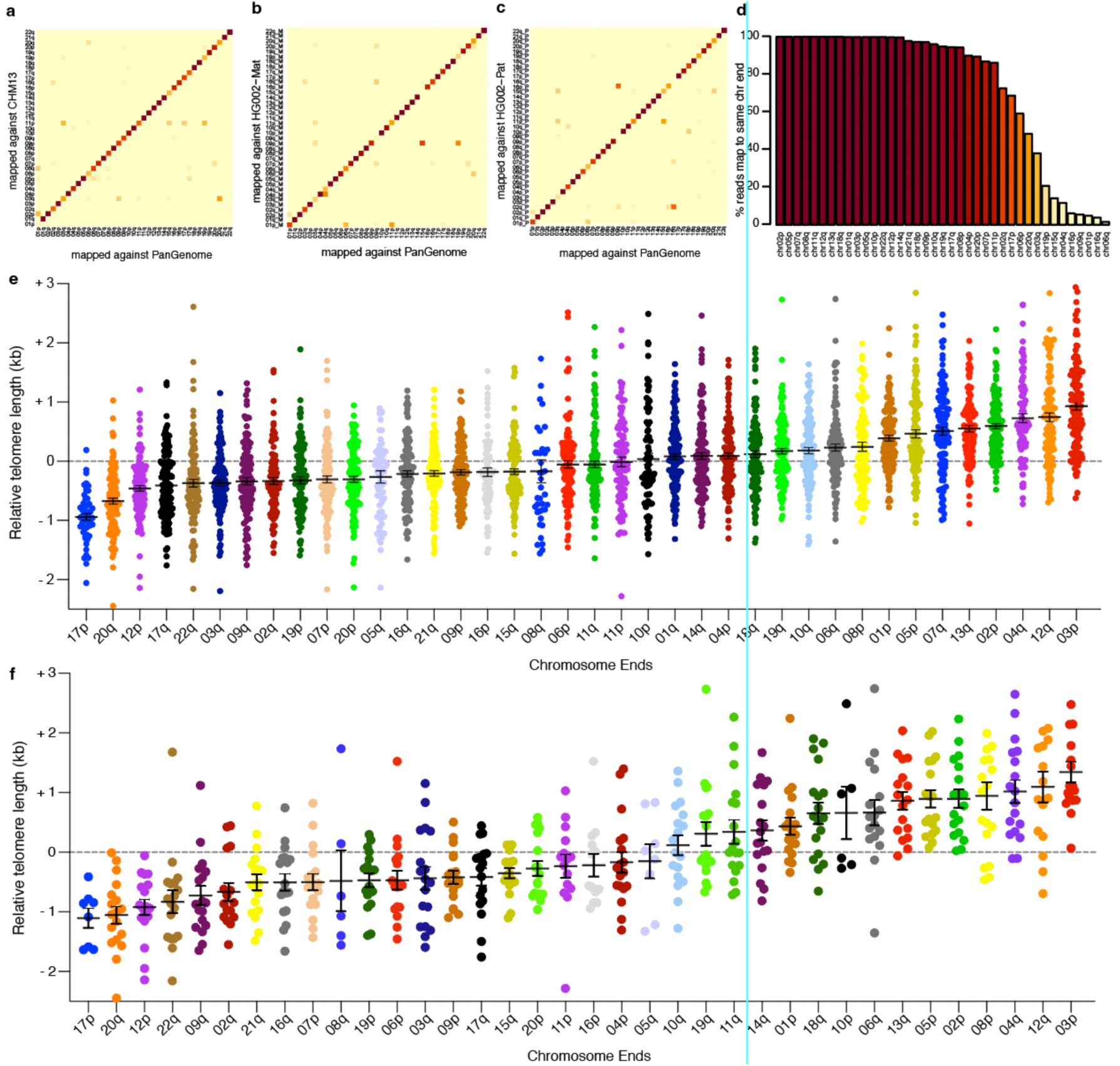
Conserved telomere lengths across 147 individuals with reads mapping to the pangenome. We used the pangenome reference to assign reads to chromosome ends for ∼920000 telomere reads obtained from 147 individuals. **a.** Matrix heatmap shows what fraction of reads that mapped to a given chromosome end in the pangenome (column) and where they map in CHM13 (rows) with a mapq of 60. Light yellow indicates 0% and dark red indicates 100% of reads mapping to the respective CHM13 chromosome end. **b**. As in **a**. but mapping to the HG002 Maternal reference **c.** As in **a.** but mapping reads to the HG002 Paternal reference genome. **d.** Bar graph showing the fraction of reads that mapped for each chromosome end in the pangenome to the same chromosome end in all three haploid genomes (CHM13, HG002 maternal and HG002 paternal). Colors are the same as in the heatmaps **a**. **b**. and **c**. **e**. To determine the relative telomere length, we calculated grand mean telomere length for a given individual and subtracted it from the chromosome specific mean telomere length for each chromosome end in that individual. Zero indicates no difference between the specific chromosome end mean telomere length and the individual’s grand mean telomere length. Bars represent mean length of a given telomere in all individuals and whiskers represent the standard error of the mean. **f**. Same as in **e**. but for cord blood samples only.

To compare the telomere length of each chromosome end across the aging population, we established the relative mean telomere length. For the ∼640000 reads that mapped to the pangenome, we calculated the grand mean telomere length for a given individual and subtracted it from the chromosome specific mean telomere length for each chromosome end for a given individual. Zero indicates no difference between the specific chromosome end mean telomere length and the individual’s grand mean telomere length (Fig 4e). We ranked the chromosome ends by their relative telomere lengths and found that 17p, 20q and 12p tended to be the shortest telomeres in the population while 4q, 12q and 3p tended to be the longest (Fig. 4e). Thus, while haplotype specific differences in telomere length are seen in a single individual (Fig. 3a), across a population, on average, certain chromosome ends are more likely to be shorter while others are more likely to be longer than the grand mean. Remarkably, previous work using qFISH to measure telomere length on metaphase spreads in 10 individuals also found 17p, 20q and 12p among the top 4 shortest and 4q, 12q and 3p among the top 8 longest ends ^21^ strengthening the conclusion that some chromosome ends are reproducibly shorter or longer than the grand mean.

To determine whether chromosome end-specific telomere lengths are present at birth, we mapped the reads from cord blood to the pangenome and calculated the relative mean telomere lengths as described above (Fig. 4f). While we had fewer cord blood samples, and therefore fewer chromosome ends met our quality filters, we found again that 17p, 20q and 12p were shorter while 4q, 12q and 3p were longer than the grand mean. This supports previous work ^45^ that suggested that telomere length at birth is maintained with age.

## Discussion

A fundamental understanding of the mechanisms that regulate telomere length is essential to develop future disease treatments. When the telomere length distribution shifts to shorter lengths, some telomeres become critically short, initiating senescence, ^46–49^ and can cause age-related degenerative disease in humans ^18^. Inherited mutations that shift to a longer equilibrium predispose people to cancer ^3,50^ and the most frequent somatic mutations in cancer increase telomerase levels and lengthen telomeres ^4,5^. Nanopore Telomere Profiling will enable the dissection of how individual telomere lengths on specific chromosomes are maintained and may play a role triggering senescence, and ultimately in disease.

### Chromosome end-specific telomere length equilibria imply new regulatory mechanisms

The predominant protein counting model for telomere length maintenance proposes that telomere proteins that bind TTAGGG repeats repress the elongation of a given telomere in *cis* ^6,51^ and longer telomeres have more repression, allowing shorter telomeres to be preferentially elongated ^8^. This model represents a robust way to maintain a length equilibrium ^52^. However, since all telomeres have the same TTAGGG repeats, the model predicts that all telomeres would be regulated around a shared equilibrium length. The demonstration of end-specific lengths indicates that other, yet unknown factors, can play a key role modifying the set point for each unique telomere length distribution.

In yeast that lack telomerase, all chromosome end-specific length distributions shortened at similar rates ^23^, suggesting telomere elongation, not shortening, is the major influence on chromosome specific length. Telomere elongation is the sum of the frequency of elongation of any given end (telomerase recruitment) and number of repeats added per elongation event (telomerase processivity). When the sum of these events, on average, equals the rates of telomere shortening, the equilibrium point is set. However, given end-specific length distributions, it is clear that this simple view does not represent the full complexity of the system. There must be factors at specific chromosome ends that regulate telomerase recruitment, processivity, or both, to establish end specific lengths. In addition, stochastic shortening such as telomere rapid deletion ^53^ or replication fork collapse ^54,55^ may play yet unknown roles in establishing telomere length equilibrium.

### Mechanisms that may influence end specific telomere length

Subtelomeric sequences are obvious candidates to regulate end-specific telomere lengths. In yeast, subtelomere DNA binding proteins can affect telomere length ^56^, although the mechanism is not yet understood. The subtelomeric TAR1 element ^57^ present in paralogy block 23 ^26,40^ was proposed to regulate telomere length, possibly through binding CTCF and regulating expression of the lncRNA, TERRA ^58–60^. Previous studies suggested that the absence of TAR1 may correlate with shorter telomeres ^26^.

However, we did not find a direct relationship of the shortest telomere with those ends described by Dubocanin *et. al.* that lack TAR1 (8q,13p,14p, 17p, 21p, 22p Xp) in our data set. Future comprehensive analysis of the subtelomere sequences adjacent to long and short telomeres will lead to new testable models for establishment of telomere length equilibria.

Epigenetic modifications of DNA or histones may influence telomere length ^61^. Human and mouse subtelomeric regions are known to be methylated at CpG sites ^62^ and experiments in mice suggest that loss of DNA methyltransferases results in shorter telomeres ^63^. Sequences in the subtelomere could recruit chromatin modifying enzymes that might influence length regulation. Subtelomere sequences may also influence other mechanisms that have been proposed to regulate telomere length including replication timing and tethering to the nuclear periphery ^64^. The availability of nanopore Telomere Profiling will allow exploration of the role of these factors in establishing telomere length equilibria.

### Chromosome end-specific length differences are present at birth and maintained as telomeres shorten with age

Telomere length is inherited from parent to child. Evidence of this comes from the genetic anticipation in Short Telomere Syndromes; short telomeres are passed down to each generation, and the severity of disease increases across generations ^65^. Similarly, in mice heterozygous for telomerase deletion, short telomeres are progressively passed down across 6 generations causing progressive severity of disease ^66^. Twin studies have also documented the inheritance of telomere length in humans ^67^.

Analysis of chromosome end-specific telomere lengths across 147 individuals showed specific telomeres tend to be the longest or shortest, supporting a previous study using qFISH on 10 individuals that identified a similar set of chromosomes as the longest and shortest ^21^. Cord blood also showed that 17p, 20q and 12p were among the shortest and 4q, 12q and 3p were among the longest ends suggesting that telomere length differences present at birth are maintained over decades. This establishment of chromosome-end specific telomere length equilibria at birth ^45^ and maintenance of the equilibria after birth leaves little room for proposed effects of life history, psychological, or environmental exposures ^68^ on telomere length. The similarity of our data with Martens *et al.* qFISH analysis is remarkable, and our method will enable future studies to explore the biological significance of this finding. We did not prospectively choose our samples to be representative of the diversity of the human population, but rather to span a wide age range. However, future studies could be powered to examine whether certain chromosome ends are consistently the shortest or longest more broadly in a diverse human population.

### Implications for human disease

Being able to accurately measure chromosome-end specific telomere length has important implications for human disease. Nanopore Telomere Profiling determines nucleotide resolution of the length distribution and can distinguish the length of specific chromosome ends unlike Southern blots, qPCR, or FlowFISH assay. Telomere profiling employs the accessible MinION instrument that can be used in-house in any research or clinical lab, with very low start-up costs, allowing for equitable access to telomere length determination methods. This method provides the opportunity to prospectively develop clinical standards analogous to those for FlowFISH and may allow clinical length measurements in samples other than blood. In addition, having a highly reproducible assay that can be easily automated will enable experimental approaches to define new regulators of telomere length. The role telomere elongation in the immortalization of cancer cells has been known since 1990 ^29,69,70^. Having a precise tool that can be automated, will allow new approaches that may exploit telomere length modulation in cancer treatment. Finally, the identification of conserved chromosome end-specific telomere lengths implies that new, undiscovered biological mechanisms influence telomere length. Nanopore Telomere Profiling will empower the field as a whole to dissect these mechanisms, leading to new discoveries in telomere biology.

## Methods

### Human subjects

Peripheral blood mononuclear cells (PMBCs) were purchased from Stem Cell Technologies, ZenBio Inc, and Precision for Medicine. Samples were chosen from the repositories based on age to span from 0 (cord blood) to 91 years. Research consent for these samples was obtained by the respective companies. Blood samples used to calibrate the assay were de-identified excess samples from Johns Hopkins Hospital, certified as exempt by the John Hopkins University School of Medicine IRB. For the Short Telomere Syndrome analysis, all subjects provided written, informed consent before enrollment in the study. Research subject were recruited from the lung transplantation clinic at the University of Pittsburgh Medical Center. All patients were diagnosed with idiopathic pulmonary fibrosis according to consensus guidelines of the American Thoracic Society and European Respiratory Societies at the time of their enrollment ^71^.

### Cell Lines

HG002 cells were cultured in RPMI 1640 media (Gibco, Cat.11875093) supplemented with 2g/L glucose 2mM L-glutamine (Glutamax, Gibco, Cat.35050061) 15% fetal bovine serum (Gibco, Cat.26140079) and 1% penicillin-streptomycin (Gibco, Cat.15140122). PBMCs were counted using the Luna II hemocytometer (VitaScientific, Cat.LGBD10029).

### Telomere Southern blot analysis

Genomic DNA was isolated using the Promega Wizard gDNA kit (Cat.A1120, Promega) and quantified by QuBit 3.0 (Thermo Fisher) using the DNA kit (Q32853; Thermo Fisher). Approximately 1 μg of genomic DNA was restricted with *Hinf*I (NEB, Cat.R0155M) and *Rsa*I (NEB, Cat.R0167L,) and resolved by 0.8% Tris-acetate-EDTA (TAE) agarose gel electrophoresis (Invitrogen, Cat.EA0375BOX). 10 ng of a 1kB Plus DNA ladder (NEB, Cat.N3200) was included as a size reference. Following denaturation (0.5 M NaOH, 1.5M NaCl) and neutralization (1.5 M NaCl, 0.5 M Tris-HCL, pH 7.4) the DNA was transferred in 10x SSC (3M NaCl, 0.35 M NaCitrate) to a Nylon membrane (GE Healthcare, Cat. RPN303B) by vacuum blotting (Boekel Scientific). The membrane was UV crosslinked (Stratagene), prehybridized in Church buffer (0.5M Na2HP04, pH7.2, 7% SDS, 1mM EDTA, 1% BSA), and hybridized overnight at 65°C using a radiolabeled telomere fragment and ladder, as previously described (Morrish and Greider 2009). The membrane was washed twice with a high salt buffer (2x SSC, 0.1% SDS) and twice with a low salt buffer (0.5X SSC, 0.1% SDS) at 65°C, exposed to a Storage Phosphor Screen (GE Healthcare), and scanned on a Storm 825 imager (GE Healthcare). The images were copied from ImageQuant TL (GE Life Sciences) to Adobe PhotoShop CS6, signal was adjusted across the image using the curves filter, and the image was saved as a .tif file.

### FlowFISH

FlowFISH was performed in the Johns Hopkins Pathology Molecular Diagnostics Laboratory as described in Alder et al. 2018 ^17^.

### Preparation of HMW DNA

A modified DNA extraction protocol was used to produce high molecular weight DNA based on the Lucigen/EpiCentre’s MasterPureTM Complete DNA and RNA Purification Kit A (Biosearch Technologies, Cat MC85200). For HG002 cell line, fresh or frozen cell pellets were osmotically lysed in presence of 150mL of Nuclei Prep Buffer (NEB, Cat.T3052) supplemented with 5.5 mL of Rnase A (NEB, Cat. T3018L) and 5.5 mL of RNase If (NEB, Cat. M0243L) per million cells for 15 seconds and mixed by flicking. For PBMC or fresh blood samples, an optional PBS wash followed by Red Blood Cell lysis step was included (10 mins at RT) prior to hypotonic lysis with Nuclei Prep Buffer (NEB, Cat.T3052) and Rnase digestion. Nuclei from 1 million cells were lysed with 300 mL of lysis buffer supplemented with 20 mL of Proteinase K (20mg/mL) (ThermoFisher, Cat. 25530049). Lysates were incubated at 50 degrees C for a minimum of 24 hours overnight with periodic vortexing at low speeds (minimum speed to achieve swirling of the solution). 150 mL of MPC Protein Precipitation Reagent solution from the Lucigen/EpiCentre’s MasterPureTM Complete DNA and RNA Purification Kit A was added to precipitate proteins followed by centrifugation at 2000 x g for 30 mins. DNA was precipitated by adding 500 mL of cold isopropanol (100%) (Supply Store, Cat.100209) and pelleted by centrifugation (2000 x g for 20 mins). DNA pellets were washed 3X with 70% ethanol and hydrated in pre-warmed (37°C) Elution buffer (Qiagen, 10 mM Tris-Cl, pH 8.5. Cat. 19086) and incubated on HulaMixer™ Sample Mixer (Thermo Fisher Scientific, Cat. 15920D) at 37°C incubator overnight at 1rpm end over end mixing.

### Annealing of TeloTags for duplex barcode assembly

TeloTags were prepared in 100μL reactions with 5mM of each of the 6 permutations of telomere splint Extended Data Fig.1A) and 30 mM of biotinylated adapter in HiFi Taq DNA Ligase Reaction Buffer (NEB, Cat. M0647S). Annealing was done by heating to 99 degrees and slowly decreasing the temperature 1°C /min in a Veriti™ 96-Well Thermal Cycler (Applied Biosystems, Cat. 4375786). After annealing, reactions were diluted 1:100 in 1x Taq buffer and kept at 4°C. The sequences of the TeloTag and splint adapter is listed in Extended Data Table 2)

### Telomere Tagging

High molecular weight genomic DNA (gDNA) was quantified using the Qubit dsDNA BR assay kit (Thermo Fisher Scientific, Cat.Q32850). A total of 40 μg of gDNA was incubated with 3μl of ClaI (NEB, Cat. R0197S) or AsiSI (NEB, Cat.R0630L) or PmeI (NEB, Cat.R0560L), or BamHI (NEB, Cat.R0136M) for 2 hours at 37°C, with gentle flicking every 20 mins. Subsequently, the enzyme was heat-inactivated at 65°C for 20 mins. Ligations were carried out using 4 μg of DNA per 50 μl reaction.

Tagging reactions were done in 50 μL volume for each reaction in a MicroAmp™ TriFlex Well PCR Reaction Plate (Applied Biosystems, Cat. A32811), with 4μl/reaction of 0.3μM duplex TeloTag adapter, 5μl/reaction of 10X HiFi Taq DNA Ligase Reaction Buffer (NEB, Cat. M0647S), and 1μl/reaction of HiFi Taq DNA Ligase (NEB, Cat. M0647S). The TeloTagging reactions were incubated for 5 mins at 65°C in a Veriti™ 96-Well Thermal Cycler (Applied Biosystems™, Cat.4375786). Ligations were done through 15 cycles of denaturing at 65°C for 1 min, followed by annealing and ligating at 45°C for 3 mins with a 15% ramp down of rate between steps.

### Telomere Enrichment and Nanopore Sequencing

For chromosome-specific telomere length measurements, we typically used 30-40 μg of DNA per sample. A standard 3 mL tube of blood or 30 million PBMC produced ∼200 ug of DNA. For bulk telomere length measurements, as little as 5-10 μg of starting gDNA was employed. All pipetting was performed using wide bore pipette tips to minimize DNA shearing, except for addition of SPRI beads where accurate volume ratios are extremely important for successful cleanups. All the Telomere Tagging reactions were pooled in DNA LoBind (Eppendorf, Cat.0030108523) tube. Cleanup and removal of excess TeloTag adapters was done using SPRI beads (Beckman Coulter, Cat. B23318) a ratio SPRI beads to DNA of 45 µL:100 µL was used. The samples were incubated with SPRI beads rotating end over end on a Hula mixer for 20 mins at 10rpm. SPRI beads were then separated using a DynaMag™-2 Magnet (Thermo Fisher Scientific, Cat. 12321D) and washed while on the magnet twice with freshly made 85% ethanol. DNA was eluted using heated (65 °C) 1X rCutsmart Buffer. The volume of elution volume was calculated to achieve 150 ng/ml final concentration based on input DNA amount. The eluting SPRI beads were incubated for 20 mins at 65°C with gently flicking every 5 mins. SPRI beads were removed using a DynaMag™-2 Magnet.

The gDNA recovery was quantified using the Qubit dsDNA BR assay kit. Tagged gDNA was enriched using Dynabeads™ MyOne™ Streptavidin C1 (Thermo Fisher Scientific, Cat. 65001). The beads were allowed to room temperature while being resuspended on a HulaMixer™ Sample Mixer (Thermo Fisher Scientific) at 3 rpm for 1h. A ratio streptavidin to DNA of 1 μg:250ng was used. The beads were washed once in Binding Buffer from Dynabeads™ kilobaseBINDER™ Kit and resuspended in equal volume binding buffer as eluted DNA volume. The beads were then added to the gDNA sample and incubated at room temp at 1 rpm on a HulaMixer™ Sample Mixer for 20 mins. Reactions can be scaled up or down as needed, though the maximum volume of beads + gDNA + binding buffer should not exceed 1.4 ml for a single 1.5 ml Protein LoBind tube (Eppendorf, Cat. 30108442). Multiple tubes can be used and pooled at the restriction enzyme digest step. After binding to streptavidin, the beads were washed using the following sequence to remove background genomic DNA: 2x kilobaseBinder wash buffer, 2x Elution buffer (Qiagen, 10 mM Tris-Cl, pH 8.5), 1x rCutsmart Buffer.

To release telomeres, the streptavidin bead-telomere complex was resuspended in 72 μl of 1X rCutsmart, 3 μl of PvuI (NEB, Cat. R3150S) PacI (NEB, Cat. R0547L) or EcoRI (NEB, Cat.R0101M), and incubated at 37°C for 30 min, with periodic gentle flicking. The sample was then heated at 65°C for 20 mins to release any bound telomeres. If multiple tubes were used, sequential rounds of digestion can be used by adding restriction enzyme to the eluted telomere solution from the first step and incubating with streptavidin-telomere beads in the second tube. Recovered tagged gDNA was quantified using the Qubit dsDNA HS assay kit. The expected recovery was approximately 0.1-0.01% of the starting gDNA sample.

Enriched telomeres were carried forward into the standard Nanopore library prep protocol from ONT. All reactions were prepared using Ligation Sequencing Kit V10 (SQK-LSK114) kits and sequenced on R9.4.1 (Oxford Nanopore Technologies, FLO-MIN106D) flow cells. Libraries were eluted in 40 ml of elution buffer (Qiagen, 10 mM Tris-Cl, pH 8.5) with optional 15 mins of incubation at 37°C to recover long molecules. Each library was split into 3 reactions. Each reaction was sequenced on a flow cell for ∼18 hours before flow cell flushing/washing using flow cell wash kit (Oxford Nanopore Technologies EXP-WSH004) and loading of the remaining fraction. Reads were collected using MinKNOW software (5.7.5) without live basecalling.

### Cost per sample for telomere enrichment and sequencing

Library construction and nanopore sequencing cost were $750 per library including $500 for flow cell. We used multiplexing strategies to lower the cost. For bulk telomere pulldown and sequencing used 10µg DNA for each sample and multiplexed 10 for a cost of ∼$80 each sample. For chromosome-specific telomere pulldown and sequencing we started with 40µg DNA and multiplexed 8 samples for a cost of ∼$140 per sample.

### Determination of telomere/subtelomere boundary position and telomere length

To determine the length of the telomere repeats, we tested two methods. One method is based on determining the junction of the telomeres and subtelomeres in the respective reference genome (CHM13, HG002 maternal and HG002 paternal) and the second method determines the subtelomere to telomere junction in every read. For method 1, to determine the junction in a reference genome, we developed a Python algorithm named TeloBP (Telomere Boundary Point). TeloBP employs a rolling window approach, scanning from the telomere into the chromosome, identifying the telomere-subtelomere junction by detecting a discontinuity in a user defined telomeric pattern. The algorithm’s default telomeric pattern is a sequence where at least 50% of nucleotides are “GGG”. As the window moves along at six nucleotide intervals, it scores telomere similarity in 100-nucleotide segments. Variants of the telomere repeats known to be in the subtelomere do not significantly change sequence content. The junction is defined when the sequence content no longer matches a telomere like sequence content. This is calculated by averaging the similarity of a sequence with a 500 bp window, marking the start of a 50% deviation, then scanning until the increase in discontinuity plateaus, marking the subtelomere boundary. After reads are mapped to the reference genome, for each read the telomere length is determined as the number of base pairs from subtelomere junction in the reference to the TeloTag. This method incorporated many variant telomere repeats into the telomere that are not incorporated by identifying the boundary as 4X TTAGGG (Extended Data Fig 3).

In the second method we determined the subtelomeres/telomeres boundary in each read. We developed a version of TeloBP that considers common errors in the nanopore Guppy base calling. These patterns are set by default based on findings in ^27^, “[^GGG]GGG|[^AAA]AAA|TTAGG.” for G strand and “CTTCTT|CCTGG|CCC…” C strands. But the patterns can be user defined as new base callers are developed. We named this algorithm TeloNP (Telomere NanoPore). Both TeloBP and TeloNP are available on Github (https://github.com/GreiderLab).

### Custom genome for mapping telomeres

For mapping reads to the T2T genomes CHM13 and HG002 we generated custom reference genomes. We first extracted the terminal 500kb of chromosome end for each genome, then removed the telomere repeats (as determined by TeloBP) from the reference genome to allow for maximized weighing of subtelomere information for read mapping.

### Bioinformatic filtering of telomere reads

Reads were first filtered for any of the following telomere patterns [“TTAGGGTTAGGG”, “TTAAAATTAAAATTAAAA”, “CCCTCCGATA”, “TGGCCTGGCCTGGCC”] based on findings in previous literature ^27^. To identify reads with a TeloTag at the end and to demultiplex samples we performed a pairwise alignment of the 24bp barcodes with the terminal 300bp of each read using the pairwise Alignment function in the Bio Strings package of Bioconductor (doi:10.18129/B9.bioc.Biostrings , R package version 2.68.1, https://bioconductor.org/packages/Biostrings). The alignment score cutoff was set so the false discovery rate for our nanopore reads was < 1% based on random 24bp barcode sequences and unused ONT barcode sequences. We used Minimap2 with the -x map-ont option to map our reads to the custom genomes HG002 and CHM13 ^42^. We only considered primary alignments that started within 1 kb of the subtelomere boundary.

### Peak calling to measure telomere length with TeloPeakCounter

To examine whether the Guppy (v 6.5.7) and Dorado (v 0.3.1) base caller accuracy call the telomere length in correctly, we developed an algorithm, TeloPeakCounter, to count repeated peaks, or waves, in the electrical signal data measured by the nanopore device. These distinct repeated waves found in the telomere region of reads correspond to the TTAGGG telomere repeat sequences. TeloPeakCounter analyzes and counts these distinctive, periodic wave patterns in the electric signal data, and enables a direct measurement of telomere length. Assuming each wave represents a 6-nucleotide telomere repeat, we can compute estimated telomere lengths for a read. The code for TeloNP and TeloPeakCounter is available at GitHub (https://github.com/GreiderLab)

### Mapping HG002 subtelomere to maternal and paternal alleles

For the diploid HG002 genome some maternal and paternal subtelomere sequences are very similar and correct assignment of reads becomes difficult. We developed a two-step mapping procedure for mapping HG002 reads. In the first step, reads are mapped to the HG002 diploid genome. Mapq mapping confidence scores are set low for these mappings, as the mapper can have difficulty deciding between very similar maternal and paternal subtelomere sequences. We applied a relatively low mapq filter cutoff of 10 to the diploid mapping. In a second step we mapped the reads also to the maternal and paternal haploid genomes separately. Mapq scores generally increase for the alignments to the haploid genomes. To identify high confidence alignments, we applied a mapq cutoff of 30 to the haploid genome alignments. A read needed to map to the same chr end in both mappings and pass the two mapq cutoffs to be considered correctly assigned. There were different numbers of reads for specific chromosome ends due to the restriction enzyme sometimes cutting very near a telomere. To minimize this, we used different sets of restriction enzymes for both the initial cutting and for the release and combined the data. This allowed mapping of more reads for some chromosome ends.

### Pangenome based mapping for chromosome assignment of telomeres from diverse individuals

We mapped reads to the pangenome to efficiently capture telomere length across the diverse population. A references file of 500kb of subtelomere sequences was assembled from each of the genomes ^43^ in the pangenome. We mapped our reads from 147 individuals to this reference. We filtered for reads that had a minimum of 2kb alignment to the pangenome reference. To compare telomere length across individuals in Fig. 4, we removed acrocentric chromosomes (13p, 14p, 15p, 21p, 22p) and X and Y subtelomeres (XpYp, and Xq Yq) which recombine in the population. We added these back into the analysis for Extended Data Fig. 5.

### Statistical Analysis

To determine whether HG002 chromosome specific telomere lengths were significantly different from the individual’s grand mean telomere length, we used Analysis of the Mean (ANOM). Statistics were calculated using the R package rstatix (v0.7.0) and ANOM (v0.2) (https://cran.rproject.org/web/packages/rstatix/index.html).

## Data Availability

Data generated during the study will be made available in public sequence repository Sequence Read Archive (SRA) https://www.ncbi.nlm.nih.gov/sra and is available upon request.

## Code availability

The python code for TeloBP, TeloNP and TeloPeakCounter is available at https://github.com/GreiderLab

## Acknowledgements

We thank Dr. Rachel Green for provision of laboratory space at Johns Hopkins as well as discussions. Drs. Brendan Cormack and Christine Gao and for reviewing the manuscript. Carla Connelly, Julie Brunelle, and Margaret Strong provided experimental and logistical assistance. We thank Dr. Mary Armanios and her lab for help in the early stages of assay development and Dr. Ludmilla Danilova for statistical analysis. This work was supported by NIH grant R35CA209974 to CWG and Johns Hopkins Bloomberg Distinguished Professorship to CWG.

## Author contributions

These authors contributed to the following aspects of this work. Conceptualization CWG, KK, SS; Data curation, AG, KK, RK, RWK, AR; Formal analysis, KK, RK, AR; Funding acquisition CWG, HL; Investigation AG, VH, KK, RWK, Methodology,CWG, VH, KK, RK, RWK, AR, SS; Project administration CWG; Resources JA, CWG, JFM, KK; Software KK, RK, HL, AR ,K-TT; Supervision CWG, HL; Validation CWG, AG, KK, RK, AR; Visualization JA, KK, RK, AR; Writing original draft CWG, KK Writing – review & editing JA, CWG, AG, VH, KK, AR, SS

## Extended Data Figures

**Extended Data Fig. 1.**
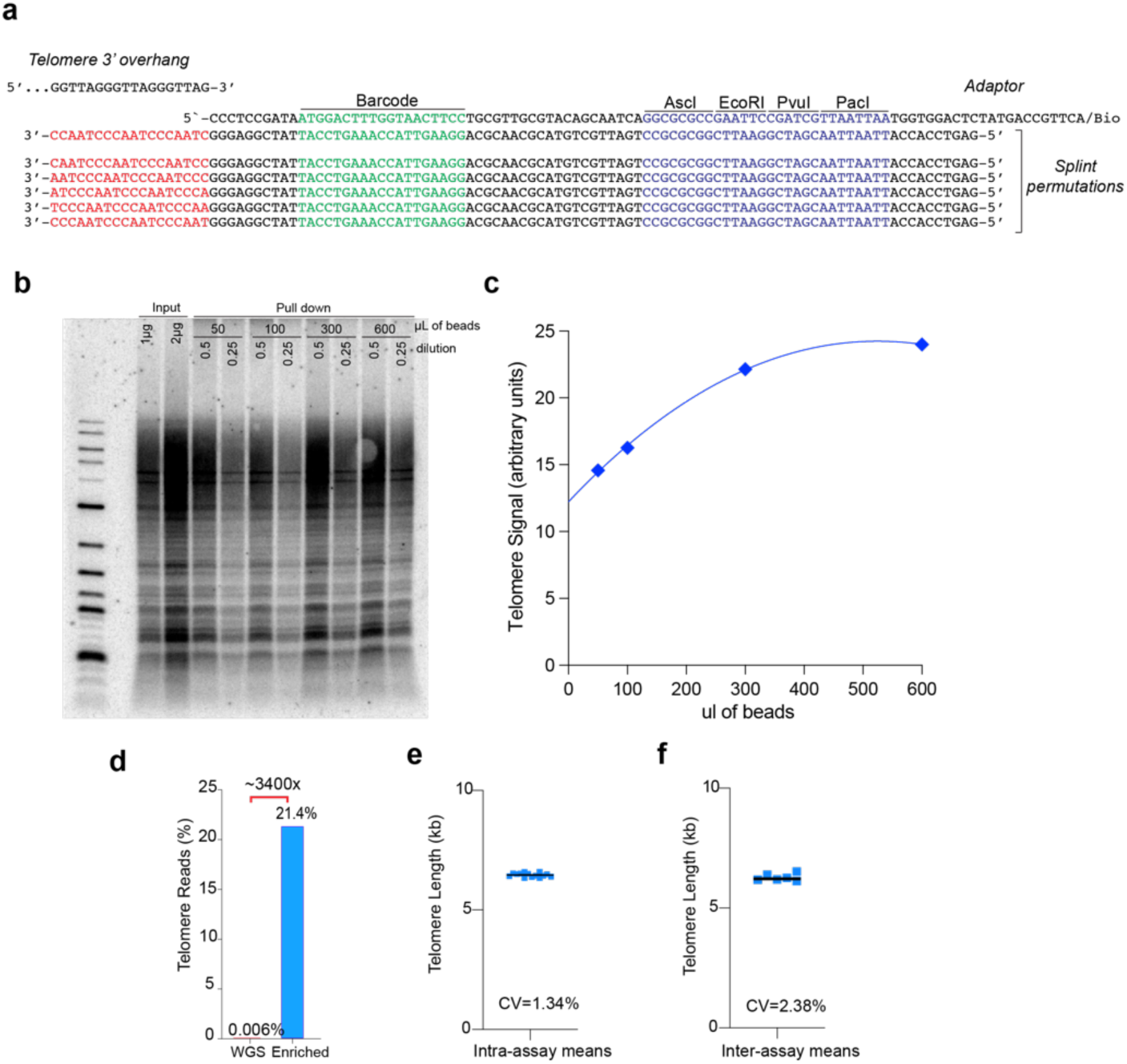
Quantitation of enrichment and assay reproducibility. **a.** Sequence of one representative TeloTag adapter. The barcoded adaptor (top strand) is annealed to a mixture of splints that have all 6 permutations of the CCCTAA sequence to improve chances of in-frame annealing to the telomere 3’ overhang. **b**. Southern blot of telomeres recovered after biotin pull down using different volumes of streptavidin bead enrichment. **c.** Quantification of the efficiency of enrichment using increasing ratio of streptavidin beads to DNA. **d.** Enrichment of telomeric reads using biotin pull down relative to WGS. **e.** Intra-assay coefficient of variation (CV) of one single with different barcodes measured multiple times on the same flow cell. **f**. Inter-assay coefficient of variation (CV) of one sample measured multiple times across different flow cells. Mean telomere length of a single sample measured on multiple different runs.

**Extended Data Fig. 2:**
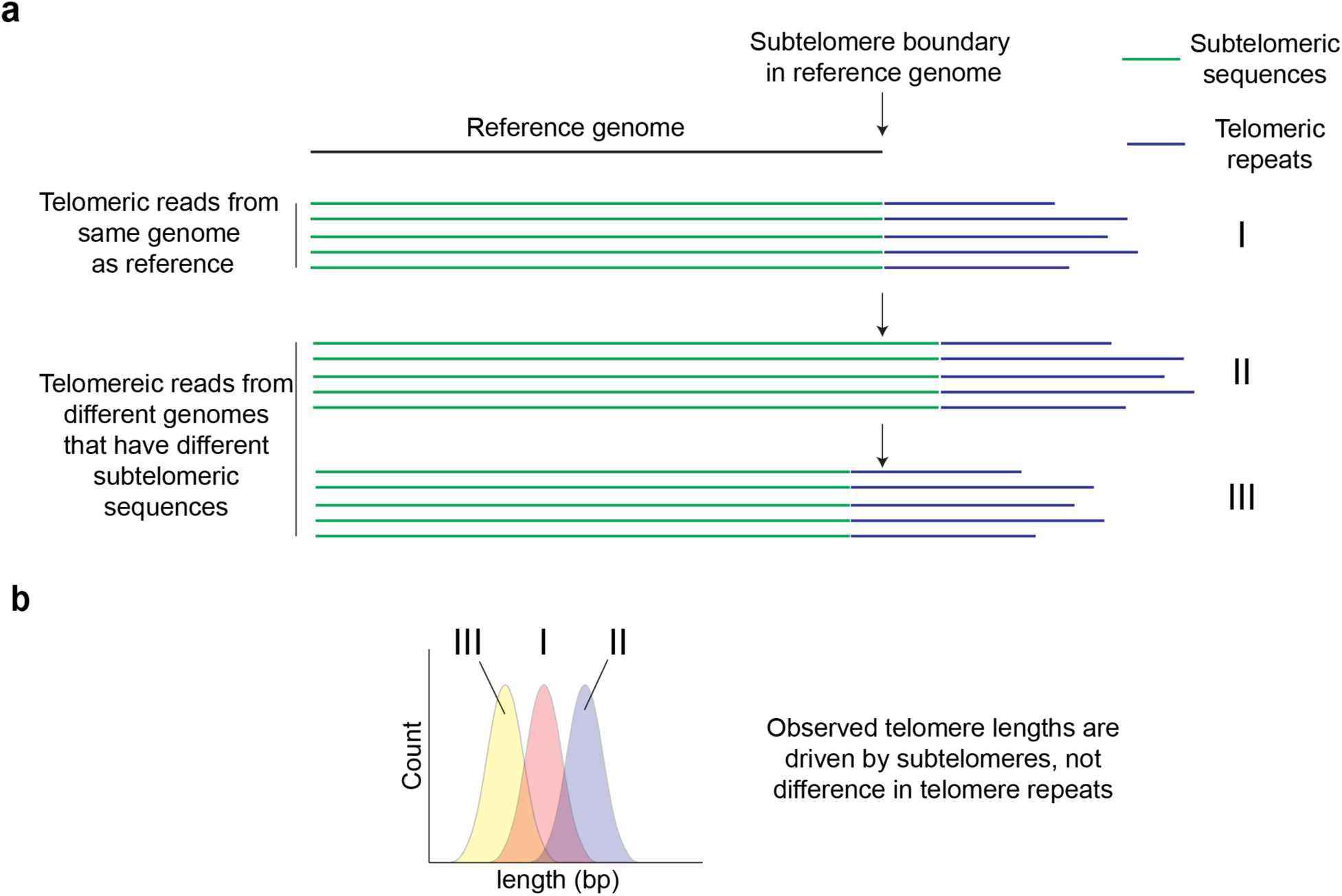
Heterogeneity in human subtelomere sequence means the telomere subtelomere boundary point can differ in sequence reads from diverse genomes and the reference genome. **a**. Telomere reads from the DNA identical to the reference genome will align at the boundary point in the reference. However, for some individuals a telomeric read will map well but there is extra sequence past the reference boundary point. For others there may be less subtelomere sequence on the read b. When telomere length is determined by mapping to the reference sequence boundary point, this can lead to incorrectly longer (II) or incorrectly shorter (III) telomere length distributions.

**Extended Data Fig. 3:**
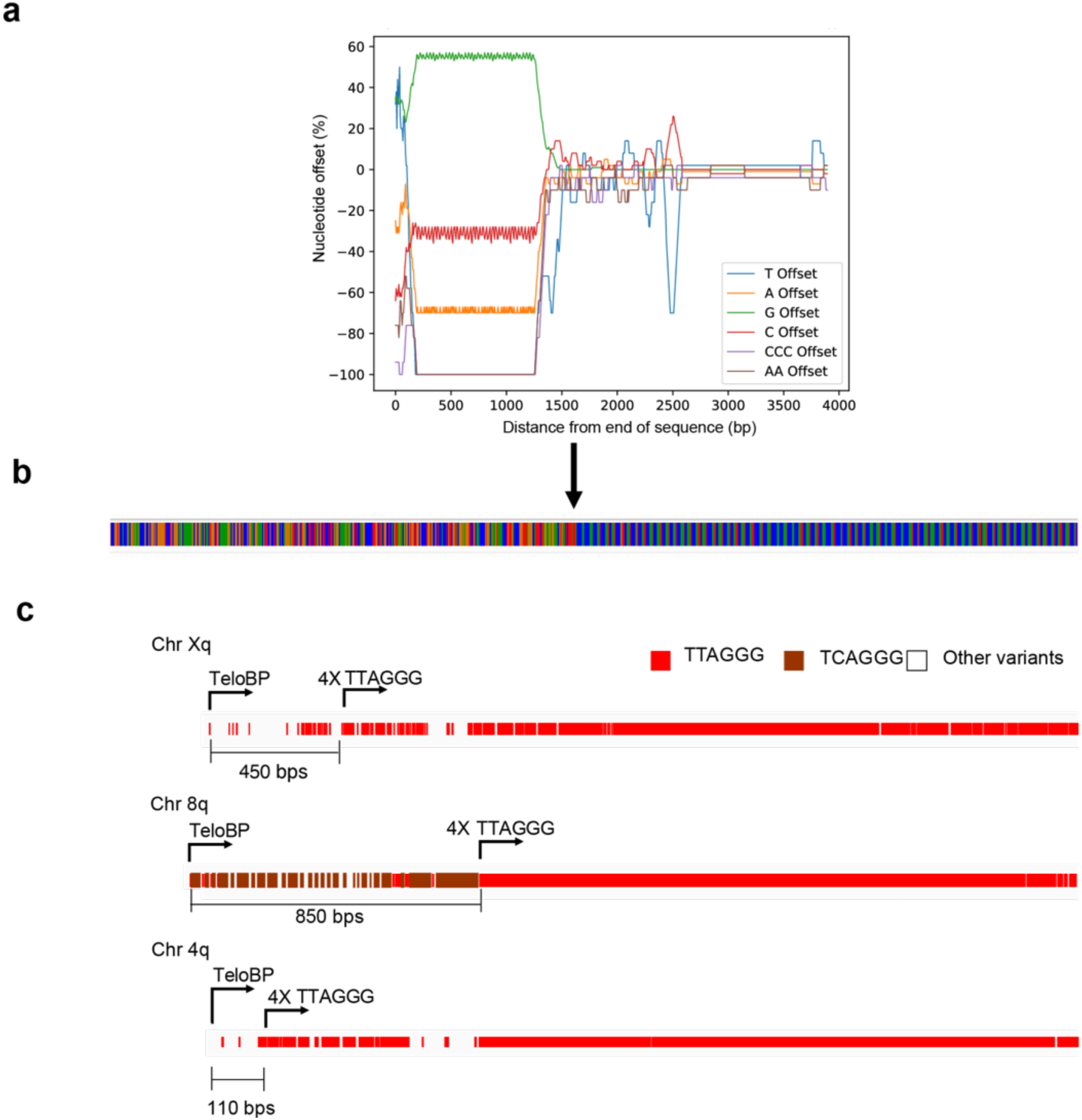
Establishing telomere boundary points with TeloBP algorithm. **a**. Representation of the nucleotide offsets for several different parameters as a rolling window scanning from telomere end on right (see methods). **b.** IGV view of the telomere sequence and where the boundary is called **c.** Example of where TeloBP incorporates variant repeats into the telomere, compared to method setting a boundary of 4 consecutive repeats of TTAGGG.

**Extended Data Fig 4:**
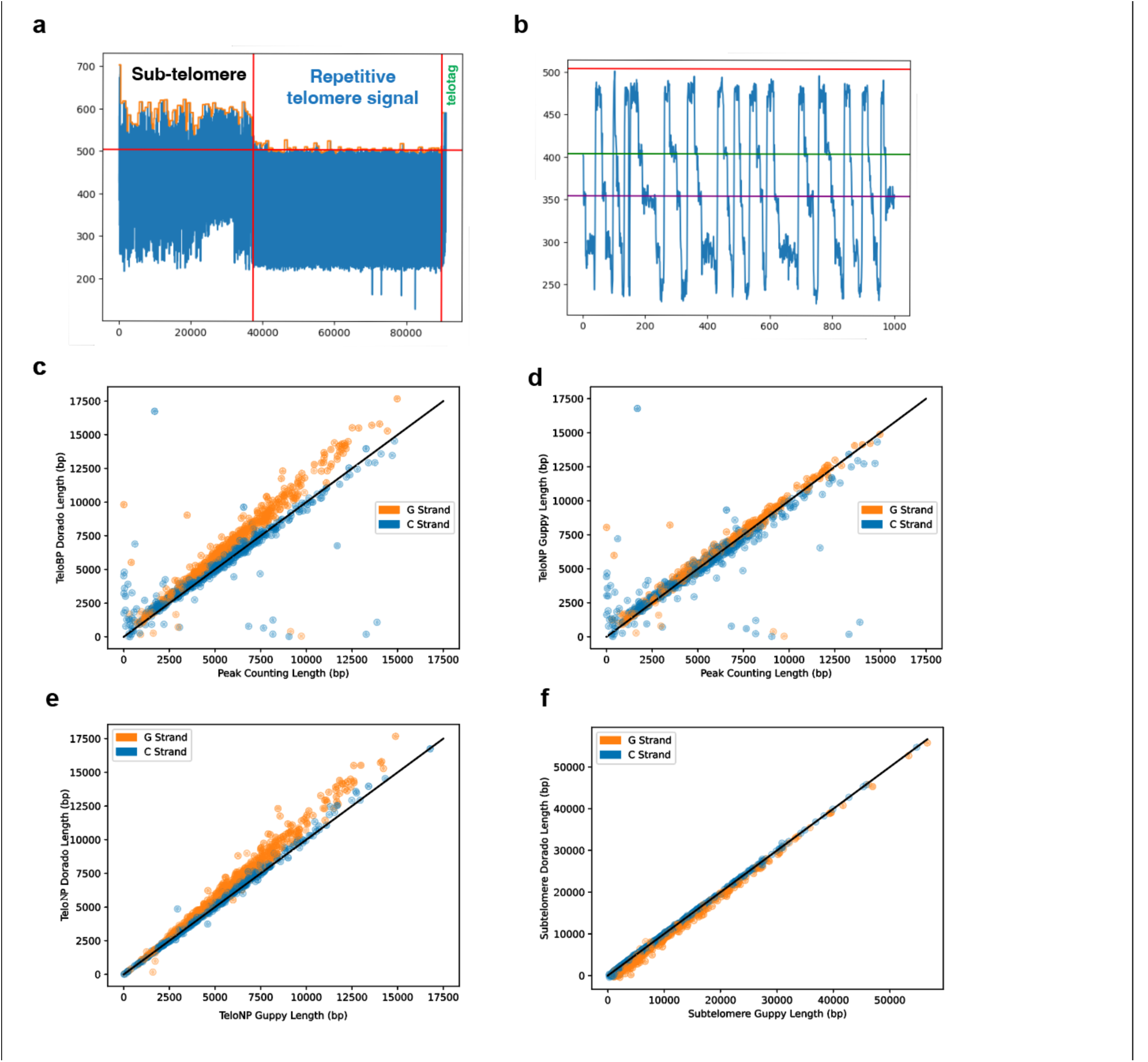
Analysis of telomere length by TeloPeakCounter. **a**. Representation of the subtelomere and telomere sequence electrical signal **b**. High resolution image of peaks in the telomere repeats in electrical signal. Comparison of Guppy (version 6.5.7+ca6d6af) versus Dorado (version 0.3.1) base caller. Each blue dot represents an individual telomere read. 2435 read were examined from one data set (F63) from Fig 2. **c**. Comparison of telomere length determined by the peak counting vs Dorado base calling Blue dots represent C-strand reads, orange dots represent G-Strand reads. **d**. Comparison of telomere length determined by the peak counting vs Guppy base calling. **e**. Comparison of Guppy telomere length by TeloNP vs Dorado. **f.** Comparison of subtelomere length with Guppy vs Dorado.

**Extended Data Fig 5.**
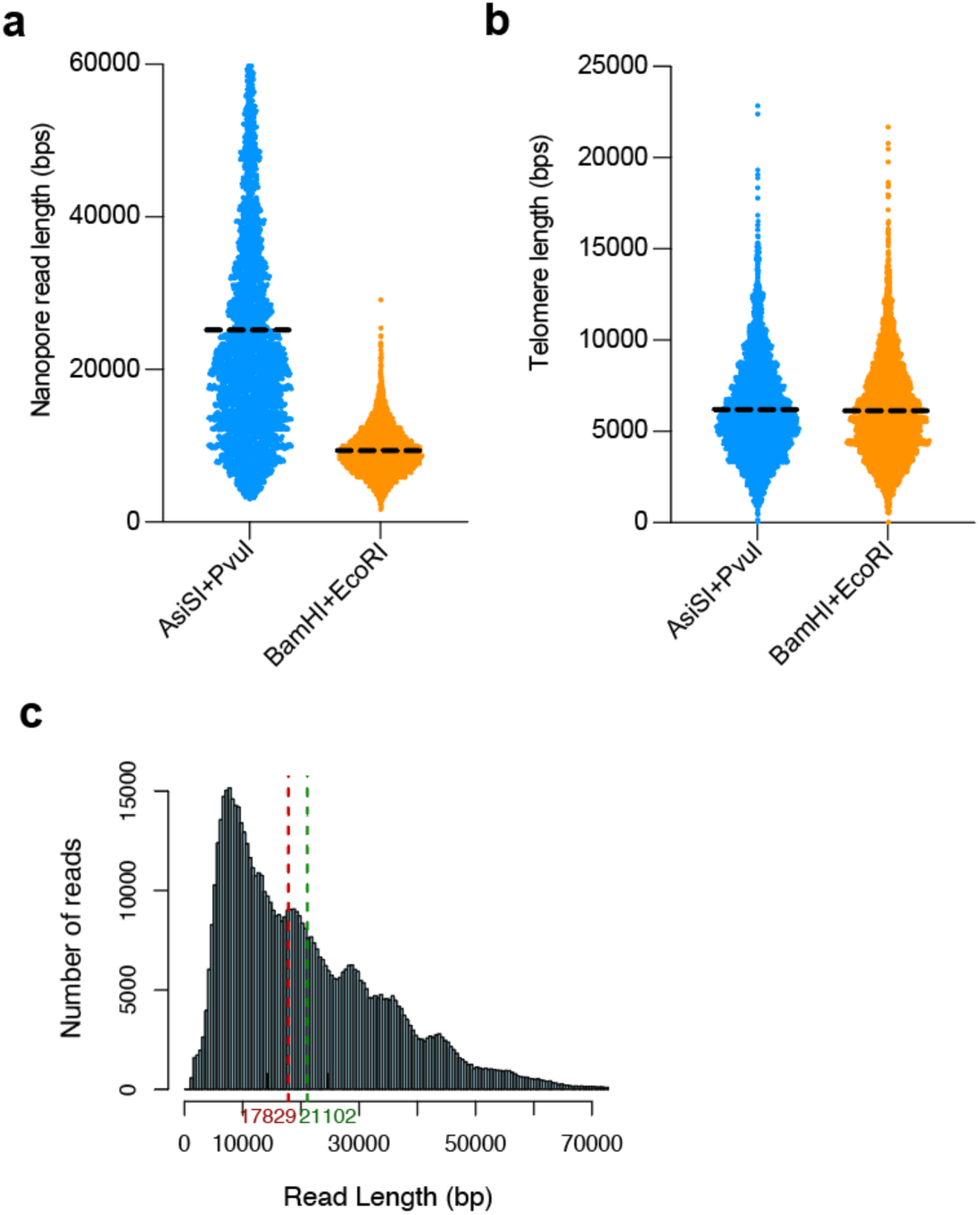
Length of fragments does not affect telomere length determination. **a.** The length of the fragments when genomic DNA is cut with AsiSI and PvuI is shown in blue. The length of the fragments when cut with BamHI and EcoRI is shown in orange. **b.** The telomere length of fragments cut with AsiSI and PvuI is in blue and BamHI and EcoRI is shown in orange. **c.** The distribution of fragment lengths for 640,000 reads that mapped to pangenome with 1kb alignment reads: the Y axis is the number of reads and the X axis is the length in base pairs The red dashed line is the median and the green dashed line is the mean length

**Extended Data Fig. 6.**
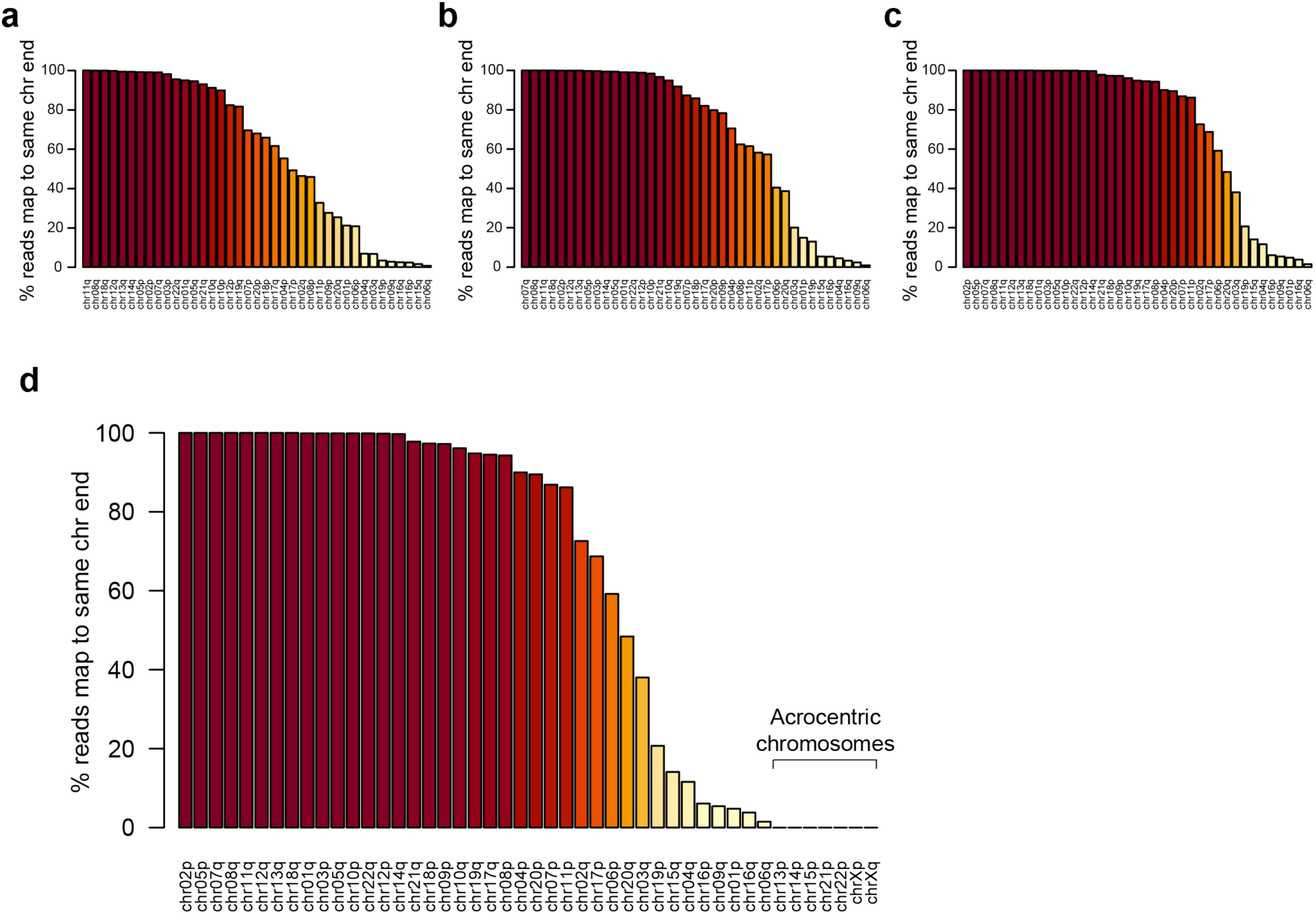
Concordance of reads mapped to the pangenome with mapping to CHM13 and HG002 Mat and HG002 Pat. We used different mapq scores to quantitate the fraction of reads that mapped to the same chromosome ends as the pangenome and the three referenced genomes **a.** Mapq score of 1 **b.** Mapq score of 30 **c.** Mapq score of 60. **d**. In previous analysis the acrocentric were omitted. Here they were included 13p,14p,15p, 21p and 22p and show less than 1% of reads mapped to the same chromosome ends for these acrocentric.

**Extended Data Table 1.**
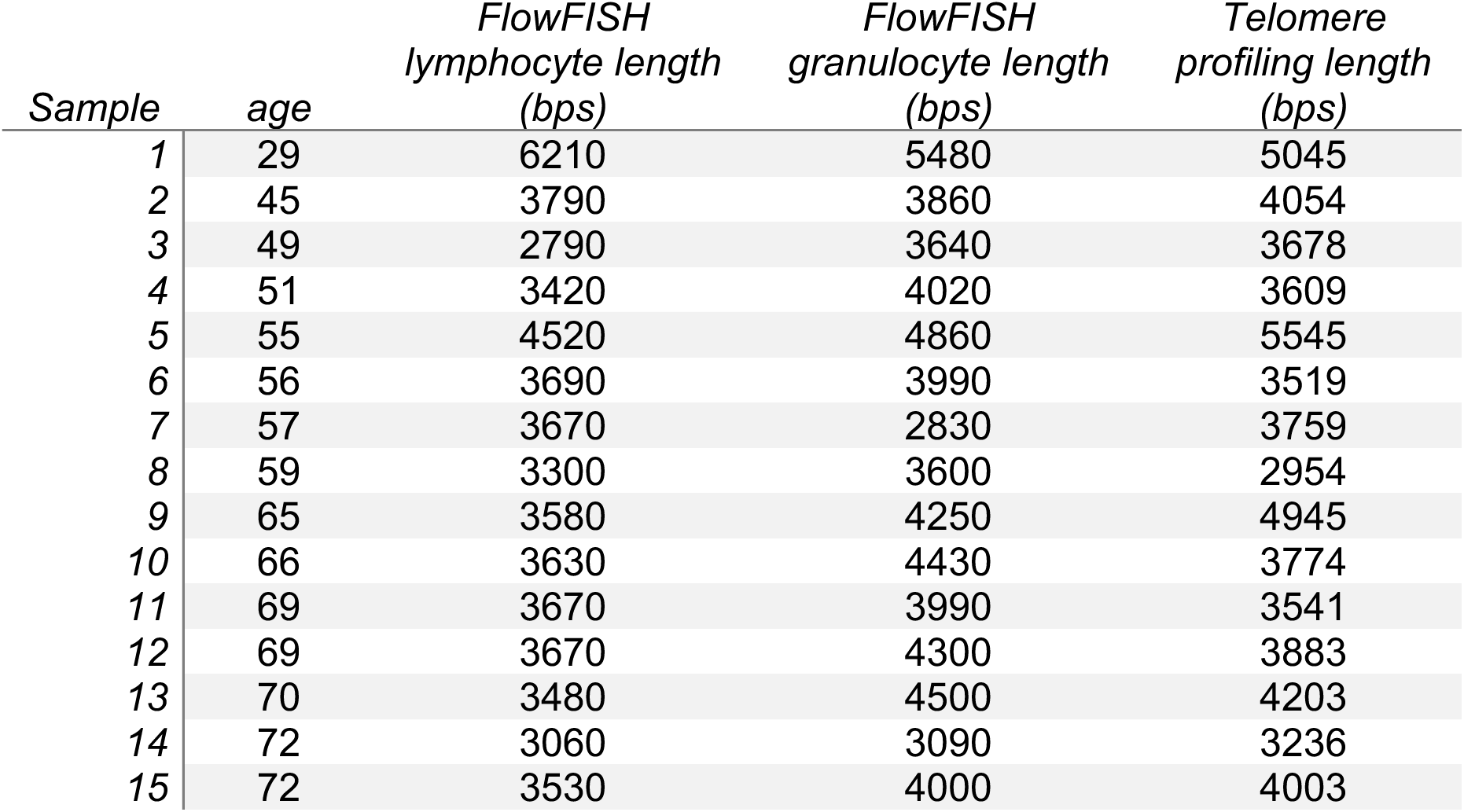
Samples included in comparison of FlowFISH and Telomere Profiling.

**Extended Data Table 2.**
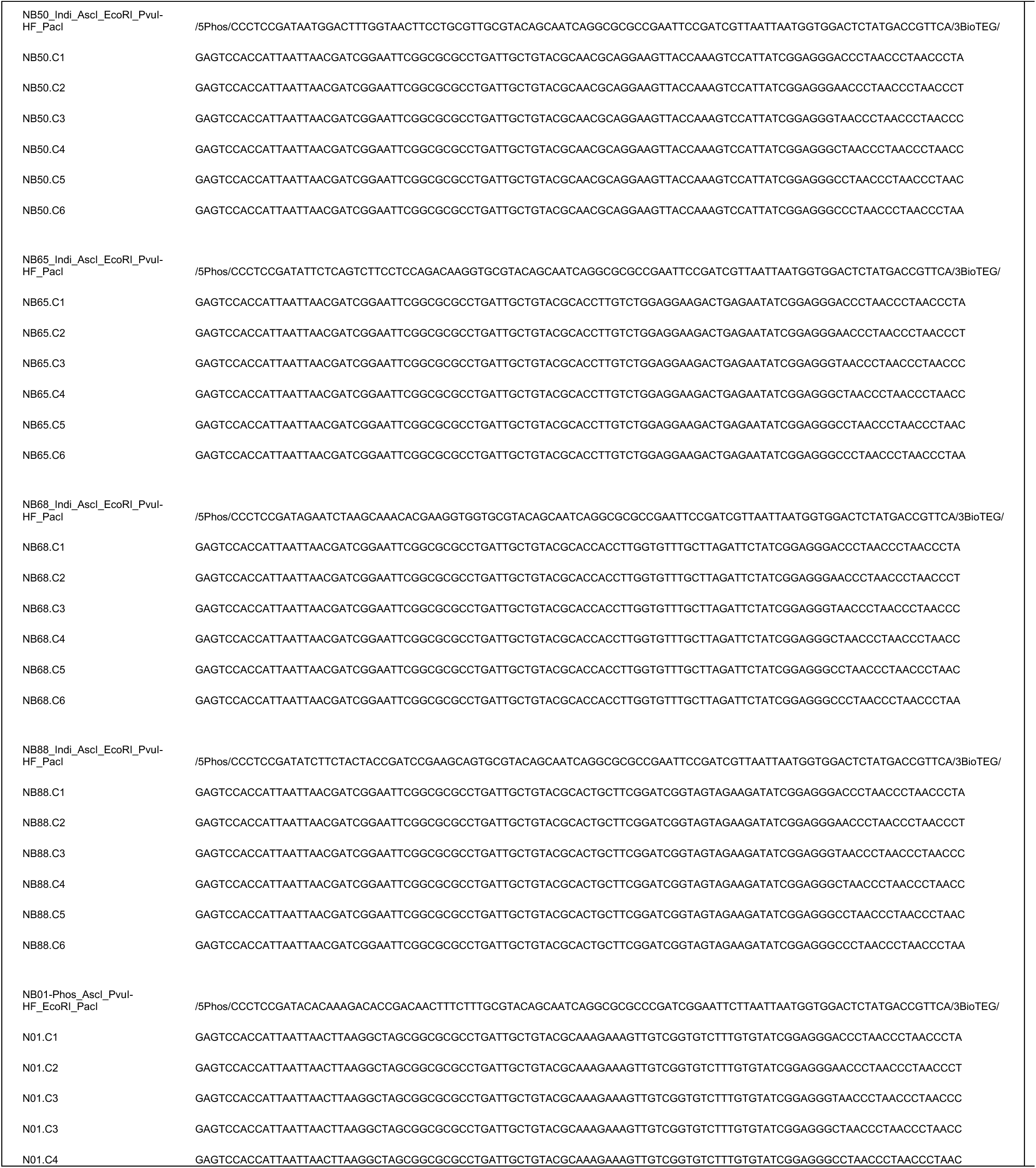
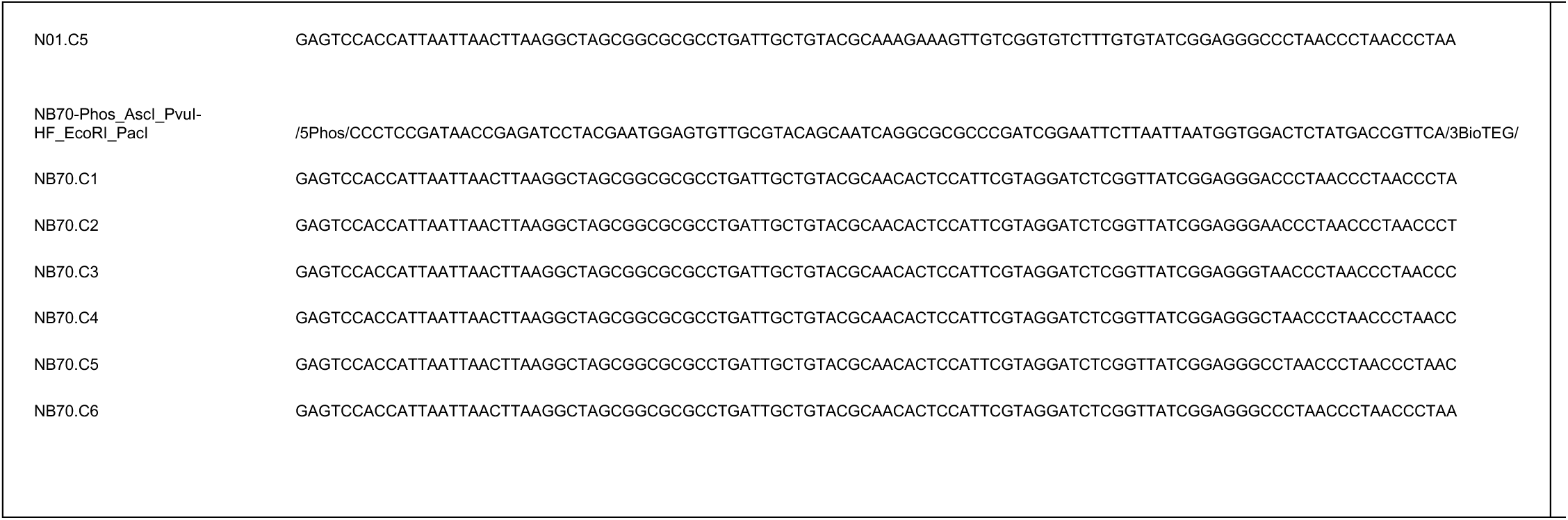
Oligonucleotides in this study.

